# Alpha-linolenic acid and dietary protein minimally but differentially regulate white adipose tissue lipolysis in female mice fed moderate-fat diets

**DOI:** 10.64898/2026.08.31.748352

**Authors:** Mathieu J Clavet, Siobhan E Woods, Melissa Gonzalez-Soto, Alexa N King, Frédéric Capel, David C Wright, David M Mutch

**Affiliations:** Department of Human Health Sciences, University of Guelph, Guelph, Ontario, N1G2W1, Canada; Unité de Nutrition Humaine, INRAE-UCA, Clermont-Ferrand, 63000, France; School of Kinesiology, University of British Columbia, Vancouver, British Columbia, Canada; BC Children’s Hospital Research Institute, Vancouver, British Columbia, Canada

**Keywords:** adipose tissue organ culture, biological sex, flaxseed, milk protein, soy protein

## Abstract

Omega-3 polyunsaturated fatty acids (n-3 PUFA) influence white adipose tissue (WAT) lipid buffering capacity; however, sex-specific regulation remains understudied. We recently reported that male mice fed a diet containing high alpha-linolenic acid (ALA) had increased WAT mass, reduced serum triglycerides and elevated lipolysis compared to mice fed a diet containing recommended levels of ALA, independent of background dietary protein. The current study examined whether subcutaneous and visceral WAT (scWAT, vWAT) lipolytic activity was altered in female C57BL/6N mice (n=16/group) fed low-ALA (1% energy) or high-ALA (3% energy) diets containing skim milk protein (SMP) or soy protein isolate (SPI) for 8 weeks. Body weight, WAT depot weights and serum triglycerides were unchanged in response to ALA content. Lipolytic markers were mostly unchanged by ALA content except for an increase in adipose triglyceride lipase (ATGL) in vWAT, while diets containing SPI modestly reduced serum cholesterol levels and increased total hormone-sensitive lipase (HSL) content in vWAT. Collectively, WAT lipolytic markers in female mice showed minimal response to diets containing high ALA content, unlike that previously reported in male mice. These results highlight the importance of considering both sexes to ensure generalizability of findings when investigating diet regulation of WAT lipid metabolism.

## INTRODUCTION

White adipose tissue (WAT) is a highly plastic and dynamic organ comprising distinct depots with both overlapping and unique functions in lipid and energy homeostasis [1]. Both subcutaneous (scWAT) and visceral (vWAT) depots play critical roles in lipid buffering, which involves the coordinated storage and release of lipids through the regulation of lipogenesis, triglyceride (TAG) synthesis, lipolysis, and fatty acid (FA) re-esterification [2,3]. Dysregulation of these processes is commonly associated with chronic overnutrition; however, emerging studies indicate that specific dietary components, independent of caloric excess, can also modulate WAT depot function [4,5].

Two dietary components, omega-3 polyunsaturated acids (n-3 PUFA) and protein, are reported to influence WAT lipid metabolism [6,7]. The three most common n-3 PUFA in the diet are the plant-derived alpha-linolenic acid (ALA) and the marine-derived eicosapentaenoic acid (EPA) and docosahexaenoic acid (DHA)[8]. These n-3 PUFA influence WAT lipid buffering capacity through a number of mechanisms, including alterations in membrane lipid composition, regulation of transcription factors and gene expression, and by serving as precursors to downstream bioactive lipid mediators [8,9]. Both EPA and DHA are understood to promote lipolysis and inhibit lipogenesis, which decreases lipid content and improves lipid utilization in WAT [10,11]. However, in an insulin-resistant state where lipolysis is upregulated, n-3 PUFA have been reported to attenuate lipolysis and limit hyperlipidemia [9,11]. Although most research to date has centered on EPA and DHA, there is growing interest regarding the independent effects that ALA may have on lipid metabolism due to recent findings using the global *Fads2^-/-^* mouse [12–14]. Moreover, studying ALA is particularly important considering that it is the predominant n-3 PUFA in the North American diet [15].

Dietary protein is also known to regulate lipid metabolism in WAT. Recent studies suggest that soy protein can decrease serum insulin levels and lipogenic gene expression in mice consuming a high-fat diet compared to mice consuming diets containing dairy proteins [16,17]. Furthermore, a previously published meta-analysis concluded that soy protein, relative to animal protein, decreased circulating cholesterol and TAG in both men and women [18]. Given the growing interest in plant-based protein products, investigations into how dietary proteins regulate WAT lipid buffering capacity is both important and timely [19].

An important caveat is that most of the prior research in the area stems from studies using male animals. This caveat is notable given that biological sex is known to influence WAT physiology and function [20,21]. For example, sexual dimorphism has been reported with respect to WAT depot distribution patterns, metabolic regulation, gene expression, hormone response, and endocrine function [21,22]. In relation to WAT lipid buffering capacity, females tend to exhibit elevated lipoprotein lipase (LPL) expression and LPL-mediated TAG hydrolysis in WAT compared to males [23,24]. Females also exhibit increased catecholamine-stimulated lipolysis and non-esterified fatty acid (NEFA) clearance from plasma, alongside reduced basal fat oxidation, relative to males [25,26]. Despite these differences, there remains a paucity of research examining responses to diet interventions in female rodent models. As such, conducting diet interventions in female animals will generate important insights to improve the generalizability of findings in the nutritional sciences.

We recently reported that a diet containing 3× the Recommended Dietary Allowance (RDA) of ALA decreased markers of WAT lipolysis in male mice, irrespective of background dietary protein (i.e., dairy versus soy)[27]. Despite aforementioned sex differences, it remains unknown whether female mice will experience similar responses to dietary ALA and background protein. Therefore, the primary objective of the present study was to examine if ALA supplementation regulates lipolytic activity in scWAT and vWAT depots in female mice. These two depots were selected due to known regional variations in lipolytic regulation [28]. Mice were fed isocaloric moderate-fat diets containing either 1% kcal from ALA, corresponding to recommended dietary intake levels, or 3% kcal from ALA, representing an elevated consumption [29]. A secondary objective was to assess whether background dietary protein (i.e., dairy or soy protein) might modulate any ALA-mediated WAT lipolytic responses.

## MATERIALS AND METHODS

### Animal Housing and Experimental Diets

Experimental plan and diet composition are identical to those previously described in Woods et al. [27], with the only difference being that female C57BL/6N mice were used in the present study. Briefly, female mice aged 10 weeks were obtained from Charles River Laboratories (QC, Canada) and individually housed in conventional ventilated shoebox cages under controlled environmental conditions (22 °C, ∼60 % humidity, 12 h light–dark cycle; ZT0 = 08:00 h) and acclimated for two weeks on standard chow (Envigo Laboratory Diets, Indianapolis, USA). Following acclimatization, mice were randomized into 4 groups (n=16 per group) and fed one of four isocaloric AIN-93G moderate-fat diets (Research Diets, NJ, USA) that varied in ALA content (1% kcal ALA [low-ALA] or 3% kcal ALA [high-ALA]) and protein source (soy protein isolate [SPI] or skim milk powder [SMP])(**Table 1**). The intervention period lasted 8 weeks, where diet and water were provided *ad libitum*. Diet macronutrient breakdown (15% kcal protein, 50% kcal carbohydrate, 35% kcal fat) was selected to align with the typical North American diet [30]. The FA composition of diets was confirmed by gas chromatography (**Table 2**). All experimental protocols were approved by the University of Guelph Animal Care Committee according to the ethical requirements of the Canadian Council of Animal Care (Animal Utilization Protocol #4350).

**Table 1.** Nutritional composition of experimental diets. Dietary composition provided by manufacturer (Research Diets Inc., NJ, USA).

| Nutritional Composition | Low ALA Milk<br>(low-SMP) |  | Low ALA Soy<br>(low-SPI) |  | High ALA Milk<br>(high-SMP) |  | High ALA Soy<br>(high-SPI) |  |
| --- | --- | --- | --- | --- | --- | --- | --- | --- |
|  | % gm | % kcal | % gm | % kcal | % gm | % kcal | % gm | % kcal |
| Protein | 16 | 15 | 16 | 15 | 16 | 15 | 16 | 15 |
| Carbohydrate | 54 | 50 | 54 | 50 | 54 | 50 | 54 | 50 |
| Fat | 17 | 35 | 17 | 35 | 17 | 35 | 17 | 35 |
| kcal/g | 4.36 |  | 4.35 |  | 4.36 |  | 4.35 |  |
| Ingredient | gm | kcal | gm | kcal | gm | kcal | gm | kcal |
| Skim Milk Powder | 420 | 1512 | 0 | 0 | 420 | 1512 | 0 | 0 |
| Soy Protein, Supro 661 | 0 | 0 | 166 | 561 | 0 | 0 | 166 | 561 |
| DL-Methionine | 3 | 12 | 3 | 12 | 3 | 12 | 3 | 12 |
| Corn Starch | 36.3 | 145 | 34.5 | 138 | 36.3 | 145 | 34.5 | 138 |
| Maltodextrin 10 | 132 | 528 | 132 | 528 | 132 | 528 | 132 | 528 |
| Lactose | 0 | 0 | 217 | 868 | 0 | 0 | 217 | 868 |
| Sucrose | 107 | 428 | 107 | 428 | 107 | 428 | 107 | 428 |
| Cellulose | 50 |  | 50 |  | 50 |  | 50 |  |
| Lard | 143.7 | 1293 | 141.5 | 1273 | 122 | 1098 | 120.1 | 1081 |
| Flaxseed Oil | 8.7 | 78 | 8.6 | 78 | 30.4 | 273 | 30 | 270 |
| t-butylhydroquinone | 0.014 |  | 0.014 |  | 0.014 |  | 0.014 |  |
| Mineral mix S10022C | 3.5 |  | 3.5 |  | 3.5 |  | 3.5 |  |
| Calcium carbonate |  |  | 13.5 |  |  |  | 13.5 |  |
| Potassium citrate, 1 H<sub>2</sub>O |  |  | 16.7 |  |  |  | 16.7 |  |
| Potassium phosphate,<br>monobasic |  |  | 1.5 |  |  |  | 1.5 |  |
| Sodium phosphate,<br>monobasic |  |  | 12 |  |  |  | 12 |  |
| Sodium chloride | 0.8 |  | 0.8 |  | 0.8 |  | 0.8 |  |
| FD&C Yellow Dye #5 | 0.05 |  |  |  | 0.025 |  | 0 |  |
| FD&C Red Dye #40 |  |  | 0.05 |  |  |  | 0 |  |
| FD&C Blue Dye #1 |  |  |  |  | 0.025 |  | 0.05 |  |
| **Total** | 917.56 | 4000 | 920.16 | 4000 | 917.56 | 4000 | 920.16 | 4000 |

**Table 2.** Fatty acid composition of experimental diets. Fatty acid composition data is expressed as relative % composition.

|  | Low ALA Milk<br>(low-SMP) | Low ALA Soy<br>(low-SPI) | High ALA Milk<br>(high-SMP) | High ALA Soy<br>(high-SPI) |
| --- | --- | --- | --- | --- |
| **Relative %** |  |  |  |  |
| C 10:0 | 0.02 | 0.03 | 0.04 | 0.01 |
| C 12:0 | 0.09 | 0.07 | 0.09 | 0.05 |
| C 14:0 | 1.21 | 1.10 | 1.07 | 0.92 |
| C 15:0 | 0.05 | 0.04 | 0.05 | 0.04 |
| C 16:0 | 22.35 | 22.12 | 19.68 | 19.51 |
| C 17:0 | 0.24 | 0.23 | 0.21 | 0.21 |
| C 18:0 | 12.87 | 12.78 | 11.47 | 11.57 |
| C 20:0 | 0.20 | 0.20 | 0.18 | 0.19 |
| C 22:0 | 0.02 | 0.03 | 0.03 | 0.04 |
| C 24:0 | 0.01 | 0.01 | 0.02 | 0.02 |
| **Total SFA** | 37.06 | 36.61 | 32.84 | 32.56 |
| C 12:1 | 0.00 | 0.00 | 0.00 | 0.00 |
| C 14:1 | 0.02 | 0.01 | 0.02 | 0.01 |
| C 16:1n-7 | 1.60 | 1.57 | 1.38 | 1.33 |
| C 16:1n-9 | 0.29 | 0.29 | 0.26 | 0.25 |
| C 18:1n-7 | 1.94 | 1.93 | 1.79 | 1.76 |
| C 18:1n-9 | 35.65 | 35.70 | 33.06 | 33.64 |
| C 20:1n-9 | 0.60 | 0.61 | 0.51 | 0.53 |
| C 22:1n-9 | 0.02 | 0.04 | 0.02 | 0.04 |
| C 24:1n-9 | 0.01 | 0.01 | 0.01 | 0.01 |
| **Total MUFA** | 40.13 | 40.15 | 37.04 | 37.56 |
| C 18:2n-6 | 16.93 | 17.52 | 16.66 | 17.69 |
| C 18:3n-6 | 0.04 | 0.03 | 0.03 | 0.02 |
| C 20:2n-6 | 0.60 | 0.60 | 0.51 | 0.51 |
| C 20:3n-6 | 0.08 | 0.08 | 0.07 | 0.07 |
| C 20:4n-6 | 0.20 | 0.18 | 0.16 | 0.16 |
| C 22:4n-6 | 0.07 | 0.07 | 0.06 | 0.06 |
| C 22:5n-6 | 0.00 | 0.00 | 0.00 | 0.00 |
| **Total N-6** | 17.92 | 18.49 | 17.50 | 18.51 |
| C 18:3n-3 | 3.95 | 3.78 | 11.71 | 10.48 |
| C 20:3n-3 | 0.10 | 0.10 | 0.09 | 0.09 |
| C 20:5n-3 | 0.01 | 0.01 | 0.01 | 0.00 |
| C 22:5n-3 | 0.05 | 0.05 | 0.04 | 0.04 |
| C 22:6n-3 | 0.02 | 0.02 | 0.02 | 0.02 |
| **Total N-3** | 4.14 | 3.97 | 11.87 | 10.64 |
| **Total PUFA** | 22.06 | 22.46 | 29.37 | 29.14 |

### Glucose and Insulin Intraperitoneal Tolerance Tests

Intraperitoneal glucose (IPGTT) and insulin (IPITT) tolerance tests were conducted in mice during weeks 5 and 6 of the feeding study, respectively, as previously described [27]. Briefly, mice were fasted for 6 h beginning at ZT0 before receiving an intraperitoneal injection of D-glucose (2 g/kg body weight) or insulin (0.75 U/kg body weight). Blood glucose was measured from a tail incision at specified intervals using a handheld glucometer (Abbott Freestyle Lite, Abbott Laboratories, IL, USA). Glucose and insulin tolerance were assessed by calculating the 120-min area under the curve (AUC) and 20-min area above the curve (AAC), respectively.

### Body Weight, Food Intake and Adipose Tissue Collection

Body weight and food intake were measured weekly. Following the diet intervention period, mice were fasted for 4 h starting at ZT0, anesthetized with isoflurane, and euthanized by decapitation post cardiac puncture for blood collection. Blood was allowed to clot for 30 min at room temperature and centrifuged for 15 min at 4°C at 1000 × g to isolate serum. Subcutaneous (scWAT) and visceral (vWAT) WAT depots were harvested for either adipose tissue organ culture or flash frozen in liquid N_2_. Serum and tissue samples were stored at -80°C until analysis.

### Serum Analyses

TAG, NEFA, glycerol, glucose and total cholesterol were measured in fasted serum samples using a Thermo Scientific Konelab clinical chemistry analyzer, according to manufacturer’s instructions.

### Adipose Tissue Lipid Extraction and Fatty Acid Composition

Adipose tissue lipids were extracted and fatty acid composition analyzed as previously described [27]. In short, adipose tissue samples (20 mg) were homogenized in 1 mL 0.1M KCl, vortexed with 4 mL 2:1 chloroform:methanol, flushed with N_2_ gas, and kept at 4°C overnight. Homogenates were centrifuged (350 × g, 10 min, room temperature) to separate the phases, where the chloroform layer was transferred to glass culture tubes, dried under N_2_, and reconstituted in 100 μL chloroform. Fractions were separated on silica TLC plates (20×20 cm; Analtech Inc, USA) with a solvent system of 80:20:1 (v/v/v) petroleum ether, ethyl ether, and acetic acid. Bands corresponding to TAG and phospholipid (PL) were visualized with ANSA under UV light, scraped, and methylated in 14% BF_3_-MeOH at 100°C for 1.5 h. Fatty acid methyl esters (FAMEs) were analyzed by gas chromatography (DB-FFAP column, Agilent Technologies, USA) and quantified using EZChrome Elite software. FA composition was expressed as a percent relative to the total FAs.

### Adipose Tissue Organ Culture and *Ex Vivo* Lipolysis

Adipose tissue organ culture (ATOC) was conducted as previously described [27]. Both scWAT and vWAT tissue (∼150 mg) were collected from a subset of mice (n=7 per diet group) and then minced, rinsed, and incubated in duplicate wells containing 2 mL lyophilized M199 media (cat#M5017, Sigma-Aldrich, ON, Canada), 50 μU/mL insulin (cat#I9278, Sigma-Aldrich) and 0.00125 μM dexamethasone (cat#D1756, Sigma-Aldrich) for 24 h at 37°C (95% O₂, 5% CO₂). Media was then replaced with fresh M199 with or without 1 μM CL 316,243 (CL; cat#C5976, Sigma-Aldrich) for 2 h. One explant per depot served as the basal control condition (i.e., unstimulated) and the other received the β-adrenergic lipolysis stimulus. After the 2-hour treatment, media and tissue were collected and stored at -80°C until analysis.

### Glycerol Quantification

Glycerol was measured in ATOC media using a commercially available colorimetric assay, as previously described [27]. Briefly, media was combined with a glycerol reagent (cat#F6428, Sigma-Aldrich) and the glycerol content in experimental samples was quantified against a glycerol standard solution (cat#G7793, Sigma-Aldrich).

### Protein Quantification and Expression Analysis

Protein extraction analysis by western blotting was done as previously described [27]. Briefly, 150 mg of tissue was homogenized (2 min, 50 m/s) in 300 μL of cell lysis buffer and then centrifuged (15 min, 4 °C at 1500 × g). The infranatant was collected and protein was quantified using a bicinchoninic acid assay against a bovine serum albumin standard, as per manufacturer instructions. Protein samples were diluted to a 1 μg/μL working concentration, and then heat denatured (5 min, 95°C). Protein samples were loaded onto 10% polyacrylamide gels and separated by gel electrophoresis (2.5 h, 90v). 10 μg protein was loaded for frozen tissue samples and 20 μg for ATOC samples. Proteins were transferred to a nitrocellulose membrane (1.5 h, 100V) which was blocked with 5% BSA in tris-buffered saline containing tween (TBST) for 1 h at room temperature, then incubated overnight in primary antibodies (1:1000 dilution for tissue samples, 1:500 dilution for ATOC samples). Primary antibodies used included ATGL (cat#2439S, Cell Signaling Technology, MA, USA), total HSL (cat#4107S, Cell Signaling Technology), phospho-HSL (pHSL) Ser^563^ (cat#4139S, Cell Signaling Technology), pHSL Ser^660^ (cat#45804S, Cell Signaling Technology), pHSL Ser^565^ (cat#4137S, Cell Signaling Technology), and phospho-pKA (cat#9624S, Cell Signaling Technology). Membranes were washed (TBST, 15 min) and then incubated with an anti-rabbit HRP conjugate secondary antibody (cat#1706515, Bio-Rad Laboratories) at a 1:3000 dilution in 5% BSA in TBST (2 h, room temperature). Membranes were incubated in ECL (cat#32106, Thermo Scientific) for 5 min to visualize bands. Membranes were imaged and bands were quantified using AlphaView Software (Alpha Innotech Software, San Leandro, CA, USA). Each membrane was also stained with Ponceau S solution (cat#P7170, Sigma-Aldrich) to normalize protein loading.

### Statistical analysis

All statistical analyses were conducted using GraphPad Prism 10 Software (GraphPad Prism, CA, USA). All data is presented as mean ± standard deviation. Tissue weights, serum markers, glucose AUC, insulin AAC, western blot and gas chromatography data were analyzed by 2-way ANOVA with main factors being ALA content (low vs. high) and dietary protein (SMP vs. SPI), as well as the ALA content × dietary protein interaction. IPGTT, IPITT, and body weight curves, as well as all ATOC protein and glycerol analyses, were analyzed using a 3-way repeated measures ANOVA using time (or CL treatment), ALA content (low vs. high) and dietary protein (SMP vs. SPI) as main factors. The main factor of time was considered for GTT, ITT, and body weight analyses, while the main factor of treatment (control vs. CL-treatment) was considered for ATOC analyses. A P < 0.05 was considered statistically significant.

## RESULTS

### ALA content and background protein did not alter body weight gain

Body weight gain in female mice did not differ between the four diet groups during the 8-week intervention **(Figure 1A).** Similarly, food intake did not differ between the four groups of mice **(Figure 1B).** Consistent with body weight, neither scWAT nor vWAT weights (normalized to body weight) were affected by the diets **(Figures 1C, D)**.

**Figure 1.**
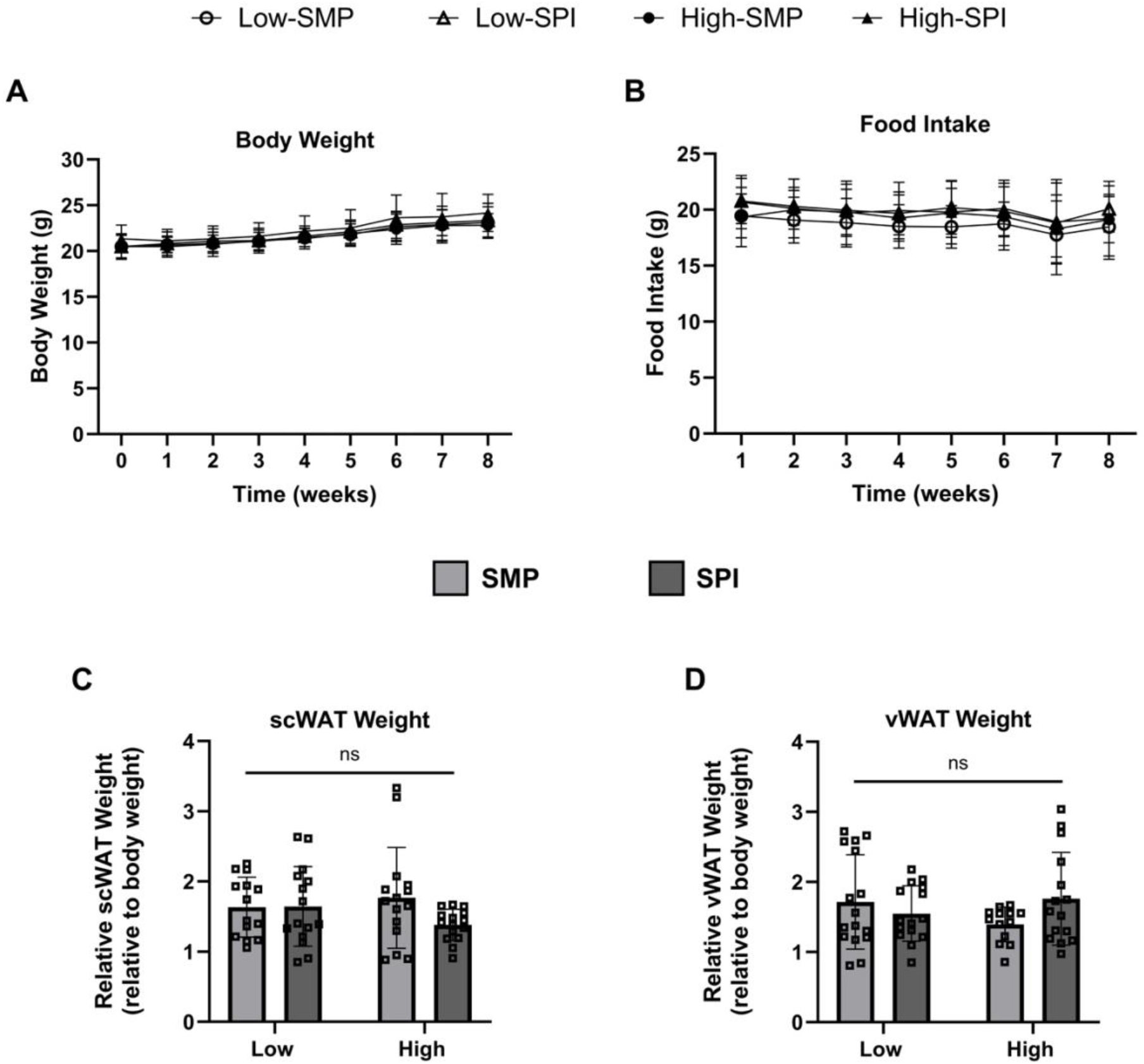
Body weight, food intake and WAT depot weights in mice fed experimental diets. Female mice (n= 14-15 per group) were fed one of four diets: 1) 1% ALA + skim milk powder (low-SMP), 2) 1% ALA + soy protein isolate (low-SPI), 3) 3% ALA + skim milk powder (high-SMP), or 4) 3% ALA + soy protein isolate (high-SPI). Body weight (A), food intake (B), relative scWAT weight (C), and relative vWAT weight (D) were measured. Body weight gain and food intake were analyzed using a 3-way repeated measures ANOVA, while WAT weights were analyzed using a 2-way ANOVA. ns = not significant. Data reported as mean ± standard deviation.

### ALA content did not modify common metabolic markers in serum, whereas soy protein decreased serum cholesterol and the NEFA/glycerol ratio

We next measured various common metabolic markers in fasted serum. Dietary ALA content did not influence serum glucose, TAG, total cholesterol, NEFA or glycerol. The NEFA-glycerol ratio, which is used as a marker of fatty acid re-esterification, was also unchanged by dietary ALA content **(Figure 2)**. In contrast, background dietary protein impacted total cholesterol levels, where mice consuming SPI-diets had lower serum cholesterol compared to mice consuming SMP-diets (P_Protein_=0.05; **Figure 2C)**. While background dietary protein did not influence NEFA or glycerol individually, the NEFA/glycerol ratio was lower in SPI-fed mice compared to SMP-fed mice (P_Protein_=0.008; **Figure 2F)**.

**Figure 2.**
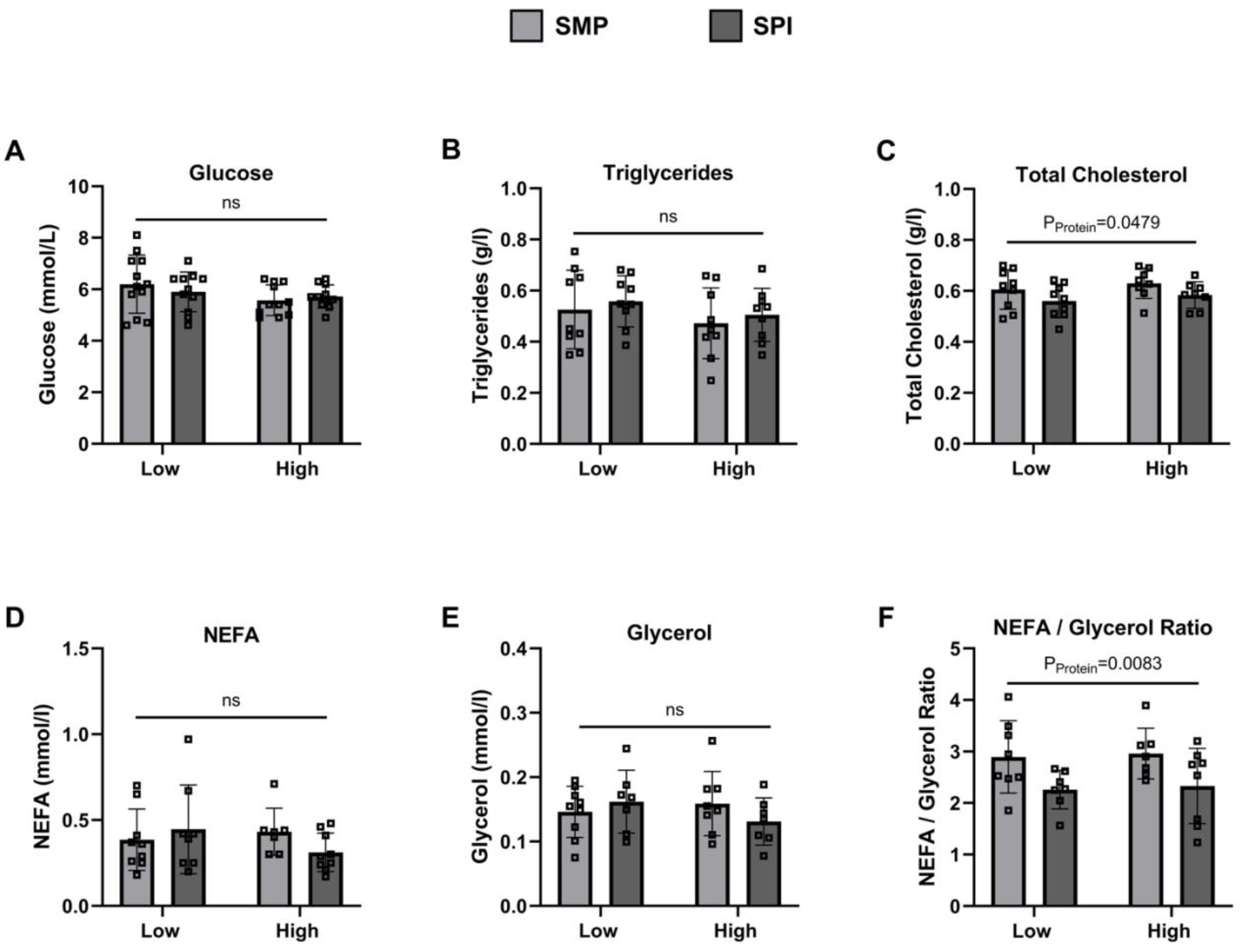
Serum biochemical measurements. Fasted glucose (A), triglycerides (B), total cholesterol (C), NEFA (D), glycerol (E), and the NEFA/glycerol ratio (F) were measured in female mice fed either low (1% en) or high (3% en) ALA in diets containing either skim milk powder (SMP) or soy protein isolate (SPI) as a source of protein. Data was analyzed using a 2-way ANOVA (n=7-11 mice per group). P_PROT_, main effect of background dietary protein, ns = not significant. Data reported as mean ± standard deviation.

### ALA content and background protein did not modify glucose and insulin tolerance

We conducted both glucose and insulin tolerance tests to assess potential changes in glucose homeostasis in response to the diets. No differences in glucose tolerance were observed between the four diet groups **(Figures 3A, C)**. Similarly, we found no effect of ALA content or dietary protein on insulin response in female mice **(Figures 3B, D)**.

**Figure 3.**
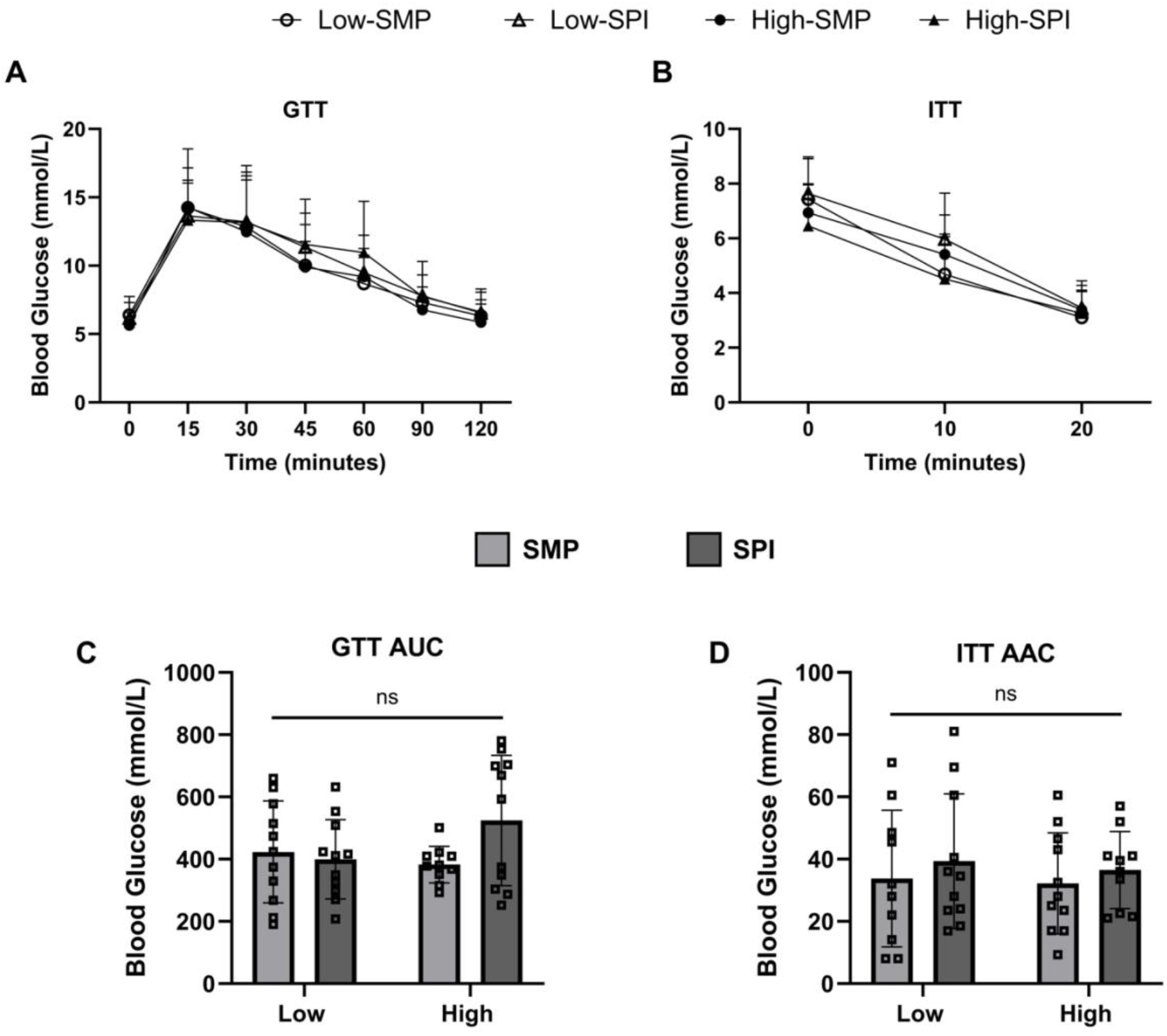
Glucose tolerance and insulin tolerance measurements. Female mice (n=11-12 per group) underwent glucose tolerance (A) and insulin tolerance (B) tests. Area under the curve (AUC) for glucose tolerance (C) and area above the curve (AAC) for insulin tolerance (D) are shown. Data was analyzed using a 2-way ANOVA and reported as mean ± standard deviation. ns = not significant.

### ALA more strongly influences TAG and PL fatty acid composition in WAT depots than background protein

We ran gas chromatography to ensure that dietary ALA content was incorporated into the two predominant lipid fractions (TAG and PL) in both scWAT and vWAT depots. WAT depot FA profiles generally reflected the FA composition of the diets. In both depots, mice provided the high-ALA diet demonstrated higher TAG-ALA levels compared to those on a low-ALA diet **(Figures 4A, 5A).** Higher TAG-ALA was accompanied by elevated TAG-EPA in both depots, and TAG-DHA in the vWAT depot **(Figures 4A, 5A).** The high-ALA diet had lower TAG-MUFA and higher TAG-n3PUFA in both depots, as well as lower TAG-SFA in vWAT **(Figures 4B, 5B).** In both scWAT and vWAT, there was no ALA response observed in TAG-n6PUFA **(Figures 4B, 5B).**

**Figure 4.**
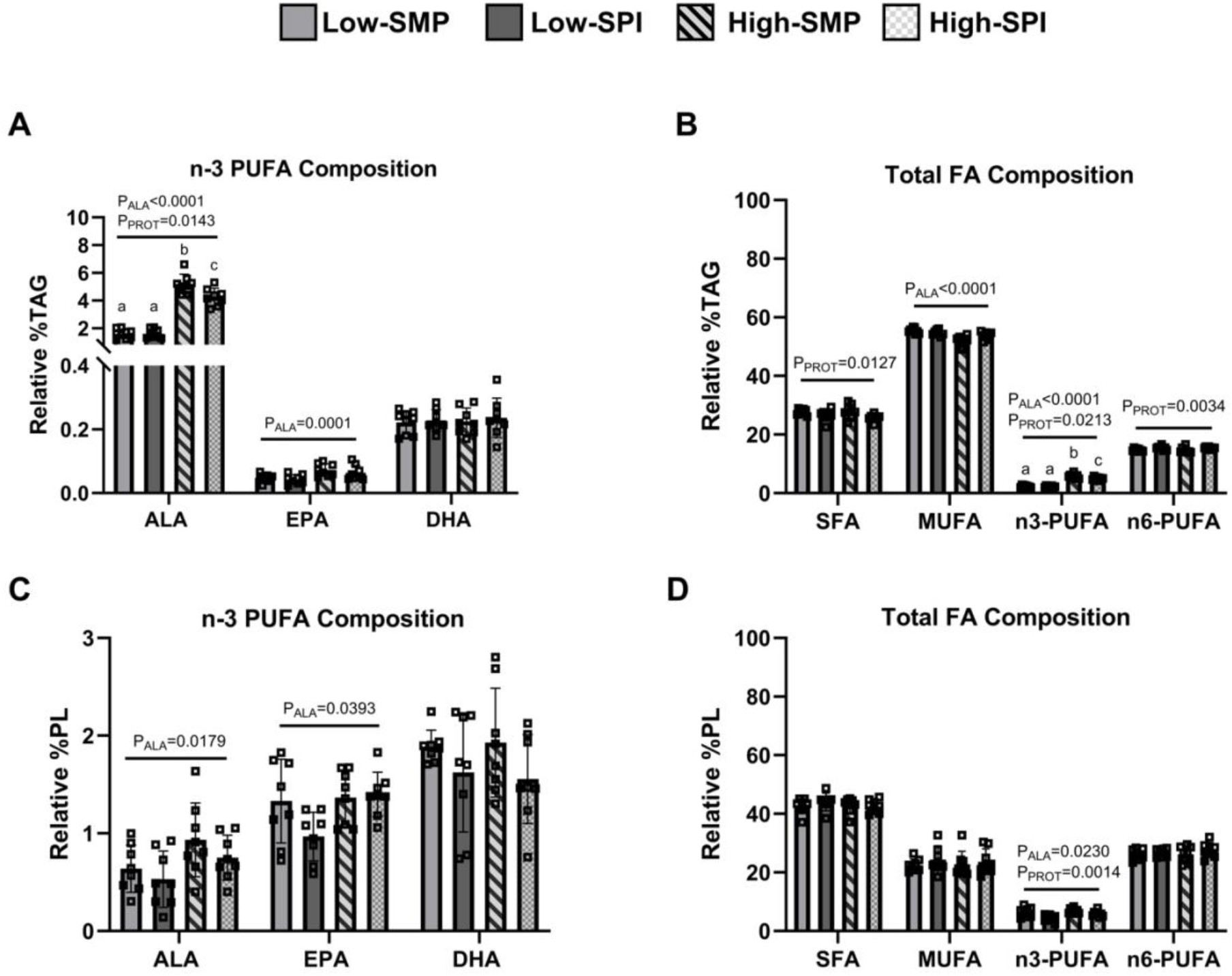
Fatty acid composition of triglycerides and phospholipids in scWAT. The relative proportions of ALA, EPA, and DHA are shown in the triglyceride (A) and phospholipid (C) fractions. The relative proportions of total saturated (SFA), monounsaturated (MUFA), n-3 polyunsaturated (n3-PUFA) and n-6 polyunsaturated (n6-PUFA) fats are also shown in the triglyceride (B) and phospholipid (D) fractions. Data was analyzed by 2-way ANOVA (n=8-9 mice per group) and reported as mean ± standard deviation. P_ALA_, main effect of dietary ALA, P_PROT_, main effect of background dietary protein. Letters above bars indicate statistically significant differences; bars with different letters are significantly different (p < 0.05).

Response to ALA was less pronounced and more variable in the PL fraction in both depots. PL-ALA and PL-EPA were higher in response to the high-ALA diet compared to the low-ALA diet, with no change in TAG-DHA levels **(Figures 4C, 5C).** In scWAT, PL-n3PUFA was higher in mice consuming the high-ALA diet; however, no response to diet was observed in PL-SFA, PL-MUFA, and PL-n6PUFA **(Figure 4D).** Mice that received the high-ALA diet also had higher PL-SFA and lower PL-MUFA in vWAT, while there was no response in both PL-n3PUFA and PL-n6PUFA **(Figure 5D).**

**Figure 5.**
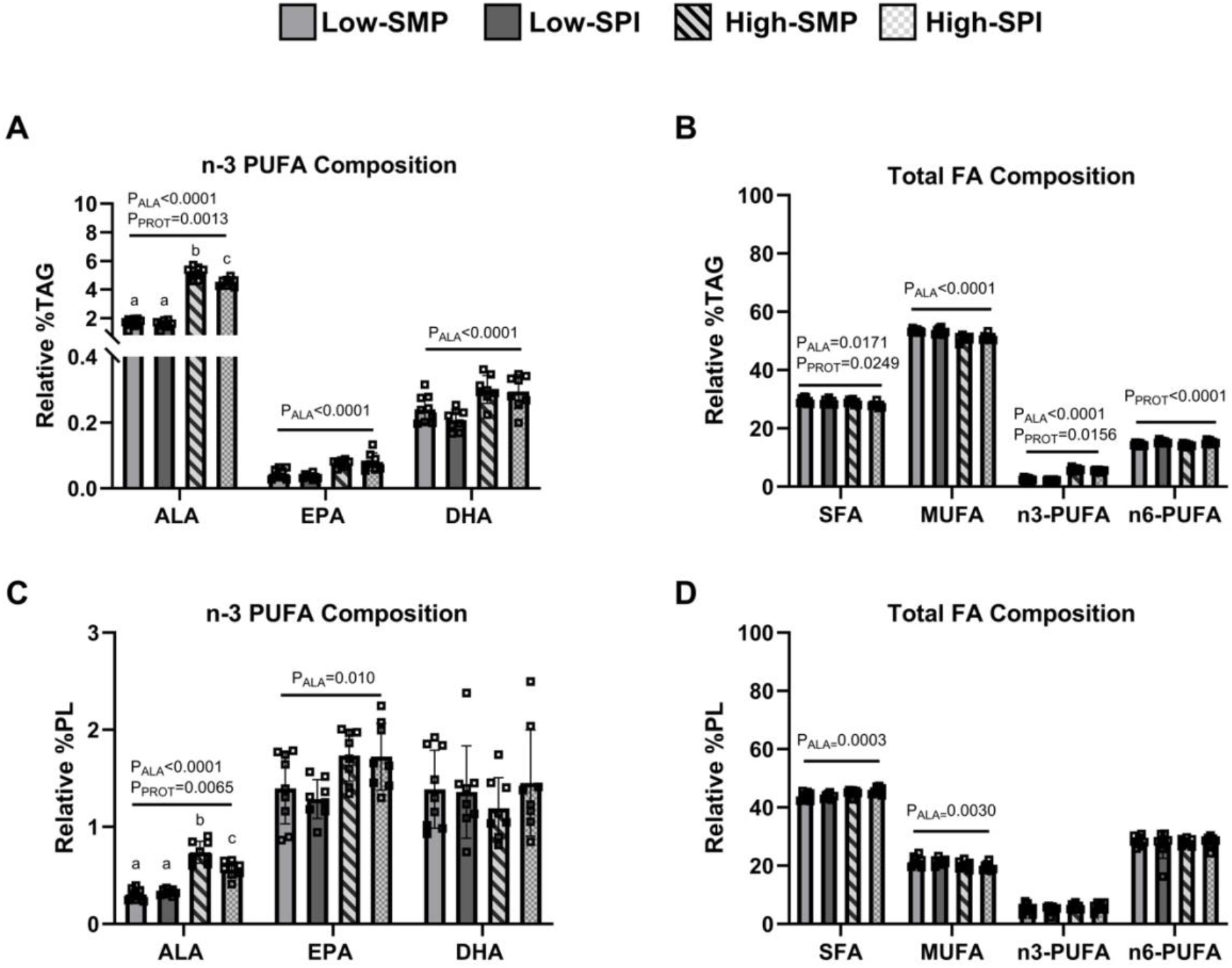
Fatty acid composition of triglycerides and phospholipids in vWAT. The relative proportions of ALA, EPA, and DHA are shown in the triglyceride (A) and phospholipid (C) fractions. The relative proportions of total saturated (SFA), monounsaturated (MUFA), n-3 polyunsaturated (n3-PUFA) and n-6 polyunsaturated (n6-PUFA) fats are also shown in the triglyceride (B) and phospholipid (D) fractions. Data was analyzed by 2-way ANOVA (n=8-9 mice per group) and reported as mean ± standard deviation. P_ALA_, main effect of dietary ALA, P_PROT_, main effect of background dietary protein. Letters above bars indicate statistically significant differences; bars with different letters are significantly different (p < 0.05).

In contrast, background dietary protein had only minimal effects on TAG and PL fatty acid composition in both depots. In scWAT, a significant ALA × Protein interaction was observed, with SPI-fed mice exhibiting lower TAG-ALA (P_ALA x Protein_=0.0152; **Figure 4A)** and TAG-n3PUFA (P_ALA x Protein_=0.0160; **Figure 4B)** compared to SMP-fed mice under high-ALA conditions. A similar interaction was identified in vWAT, where SPI-fed mice showed lower TAG-ALA (P_ALAx Protein_=0.0193; **Figure 5A)** and PL-ALA (P_ALA x Protein_=0.0007; **Figure 5C)** relative to SMP-fed mice under high-ALA conditions. Additionally, there was a main effect of protein on scWAT TAG-SFA and TAG-n6PUFA **(Figure 4B)**, as well as vWAT TAG-SFA, TAG-n3PUFA, and TAG-n6PUFA **(Figure 5B)**; however, no additional protein-related effects were observed.

### ALA and background protein modulate lipolytic protein expression in vWAT but not scWAT

Next, we examined whether ALA content and/or background protein impacted key markers of lipolysis in scWAT and vWAT. In scWAT, there was no change in ATGL, HSL, pHSL Ser^563^, pHSL Ser^660^, and pHSL Ser^565^ protein content between the different diet groups **(Figures 6A-E).** In vWAT, both ALA and background protein responses were detected (**Figure 7).** Female mice fed high-ALA diets had higher vWAT ATGL protein content compared to those fed low-ALA diets **(Figure 7A)**, regardless of background dietary protein. In contrast, mice fed the SPI-diet had higher total HSL and lower pHSL Ser^565^ protein content in vWAT compared to mice fed the SMP-diet **(Figures 7B, E)**. Neither pHSL Ser^563^ nor pHSL Ser^660^ protein content in vWAT were affected by the diets **(Figures 7C, D).**

**Figure 6.**
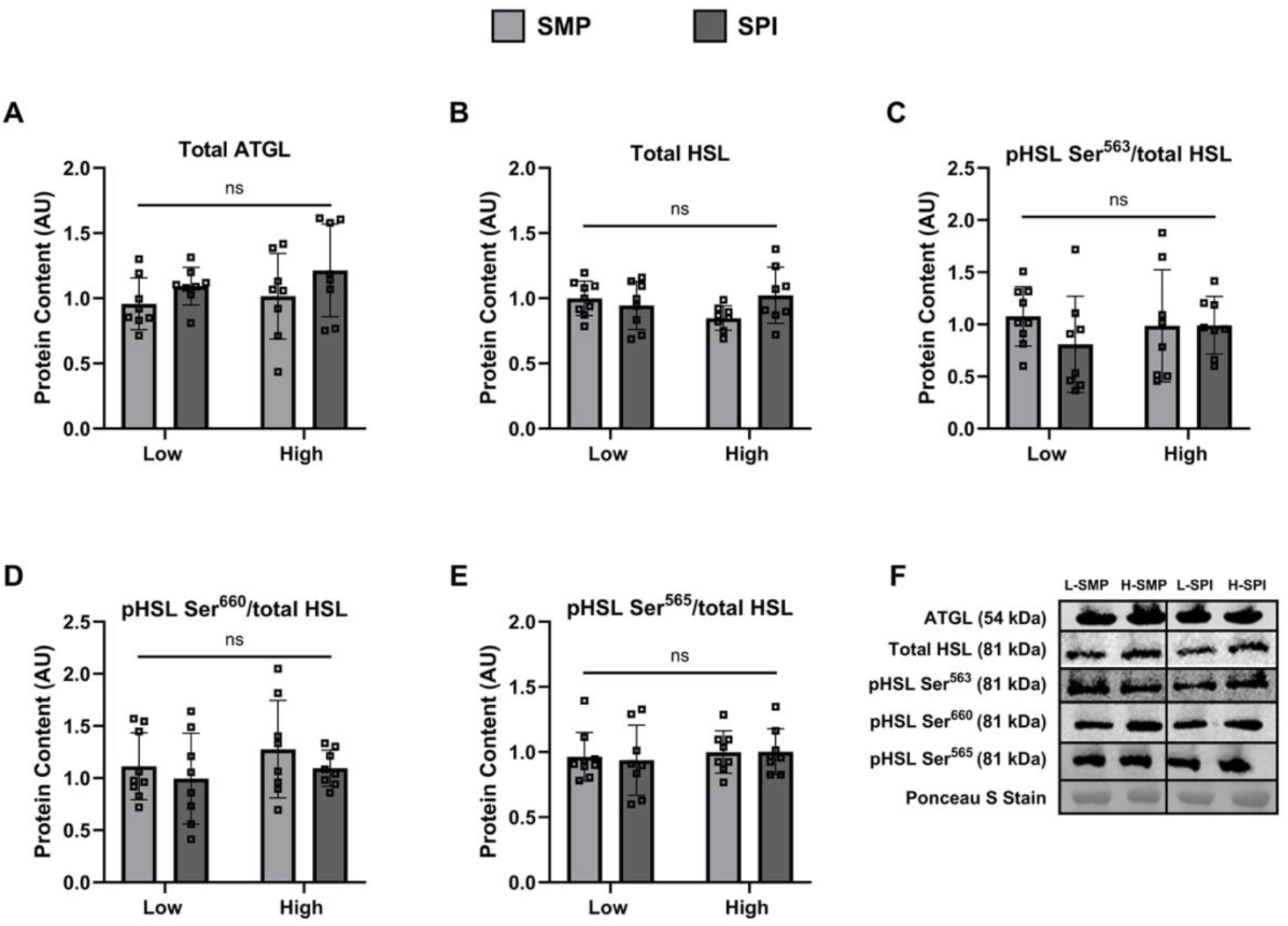
Markers of lipolysis in scWAT. The protein abundance for total ATGL (A), total HSL (B), pHSL Ser^563^ (C), pHSL Ser^660^ (D) and pHSL Ser^565^ (E) were measured in frozen scWAT samples from mice (n=8-9 mice per group). Representative blots are provided in (F). Briefly, each primary antibody was run on an individual gel and normalized to its respective Ponceau S stain to account for any differences in sample loading. Only a single representative Ponceau S stain is shown in (F). Data was analyzed by 2-way ANOVA and reported as mean ± standard deviation. ns = not significant.

**Figure 7.**
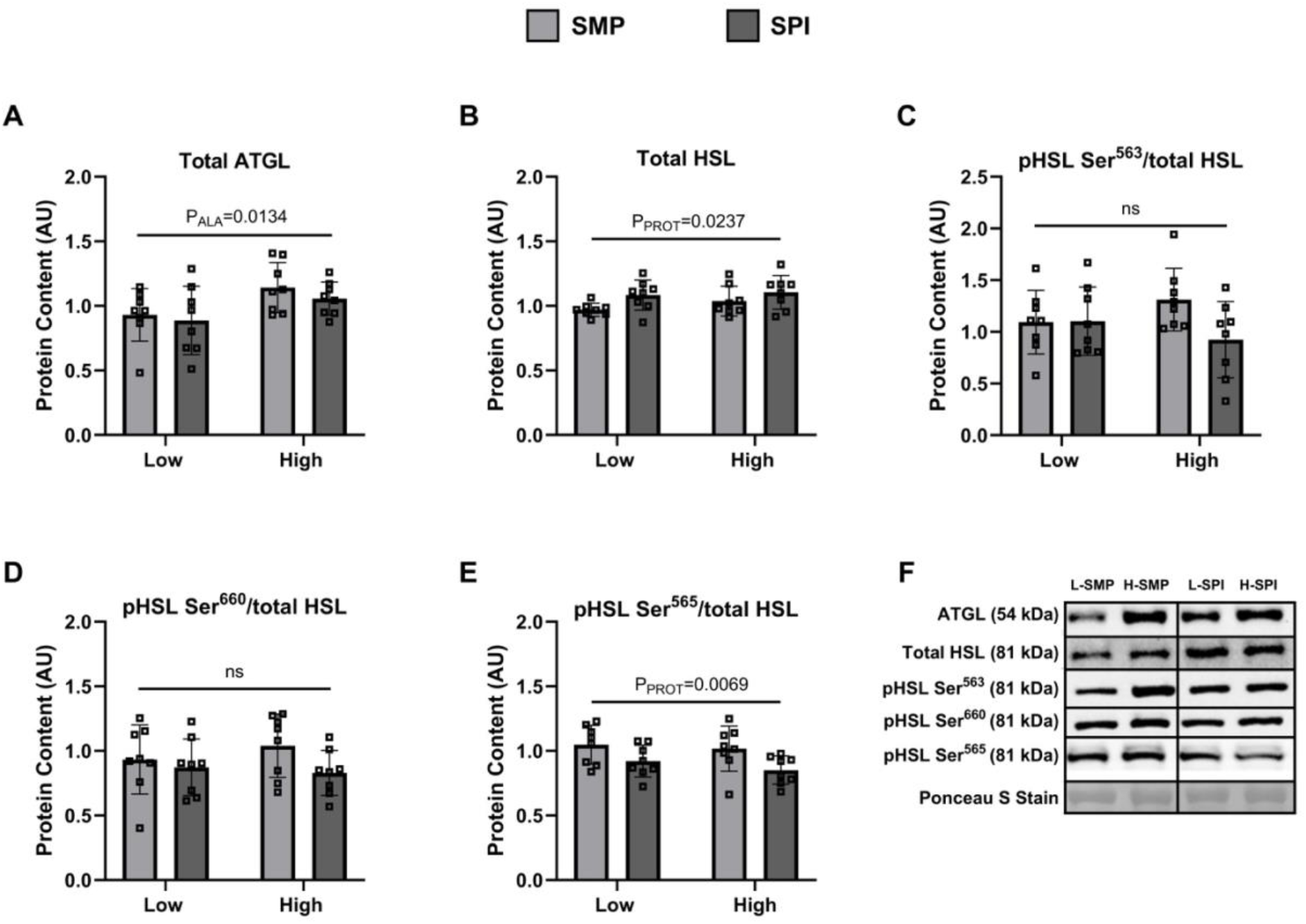
Markers of lipolysis in vWAT. The protein abundance for total ATGL (A), total HSL (B), pHSL Ser^563^ (C), pHSL Ser^660^ (D) and pHSL Ser^565^ (E) were measured in frozen vWAT samples from mice (n=8 mice per group). Representative blots are provided in (F). Briefly, each primary antibody was run on an individual gel and normalized to its respective Ponceau S stain to account for any differences in sample loading. Only a single representative Ponceau S stain is shown in (F). Data was analyzed by 2-way ANOVA and reported as mean ± standard deviation. P_ALA_, main effect of dietary ALA; P_PROT_, main effect of background dietary protein; ns = not significant.

### ALA content modulates β-adrenergic-stimulated lipolysis in cultured adipose tissue

We next conducted a functional study in cultured WAT explants treated with a β-adrenergic agonist (CL 316,243). Following a 2h treatment with CL, glycerol content in media was measured and used as a marker of lipolysis. As expected, CL treatment increased glycerol release into media from both scWAT and vWAT depots; however, neither dietary ALA nor protein differentially impacted glycerol release in the two depots in a basal state **(Figure 8)**. CL treatment did not reveal any differences in glycerol release in vWAT. Although a significant CL × ALA × Protein interaction was seen in scWAT (P_CL x ALA x Protein_=0.007; **Figure 8)**, post-hoc analyses were unable to identify the basis for this significant interaction.

**Figure 8.**
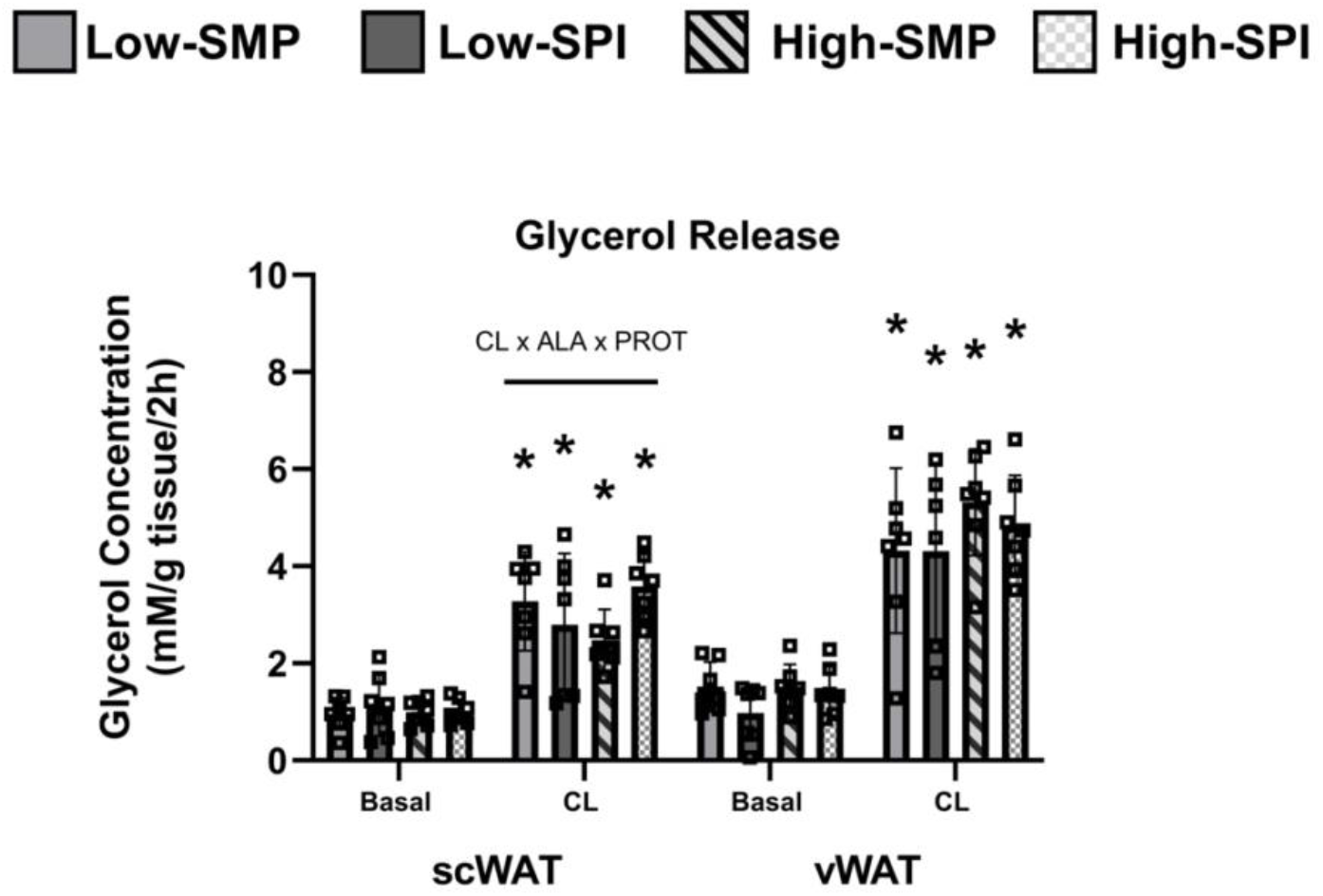
Glycerol release measured in the media of cultured scWAT and vWAT samples in basal and stimulated conditions. Adipose tissue was collected and separated for basal and stimulated conditions. β-adrenergic stimulation with CL 316,243 (CL) lasted for 2 h. * indicates significant differences between matching CL and basal conditions. CL × ALA × PROT indicates a significant interaction effect was observed in CL stimulated conditions, as determined with a 3-way ANOVA. Data reported as mean ± standard deviation. n=7 mice per group.

The lipolytic protein expression trends observed in cultured WAT explants were consistent across both scWAT and vWAT. Specifically, there was a main effect of ALA on total HSL **(Figures 9A, 10A)** and pKA substrates **(Figures 9B, 10B)**; however, post-hoc analyses did not identify significant differences between groups. Although CL stimulation significantly increased the expression of pHSL Ser^563^ **(Figures 9C, 10C)** and pHSL Ser^660^ **(Figures 9D, 10D)** in both WAT depots as expected, these increases were not differentially affected by the diets. No differences were observed at the pHSL Ser^563^ site between the four diet groups in either basal or CL-stimulated conditions (**Figures 9E, 10E)**.

**Figure 9.**
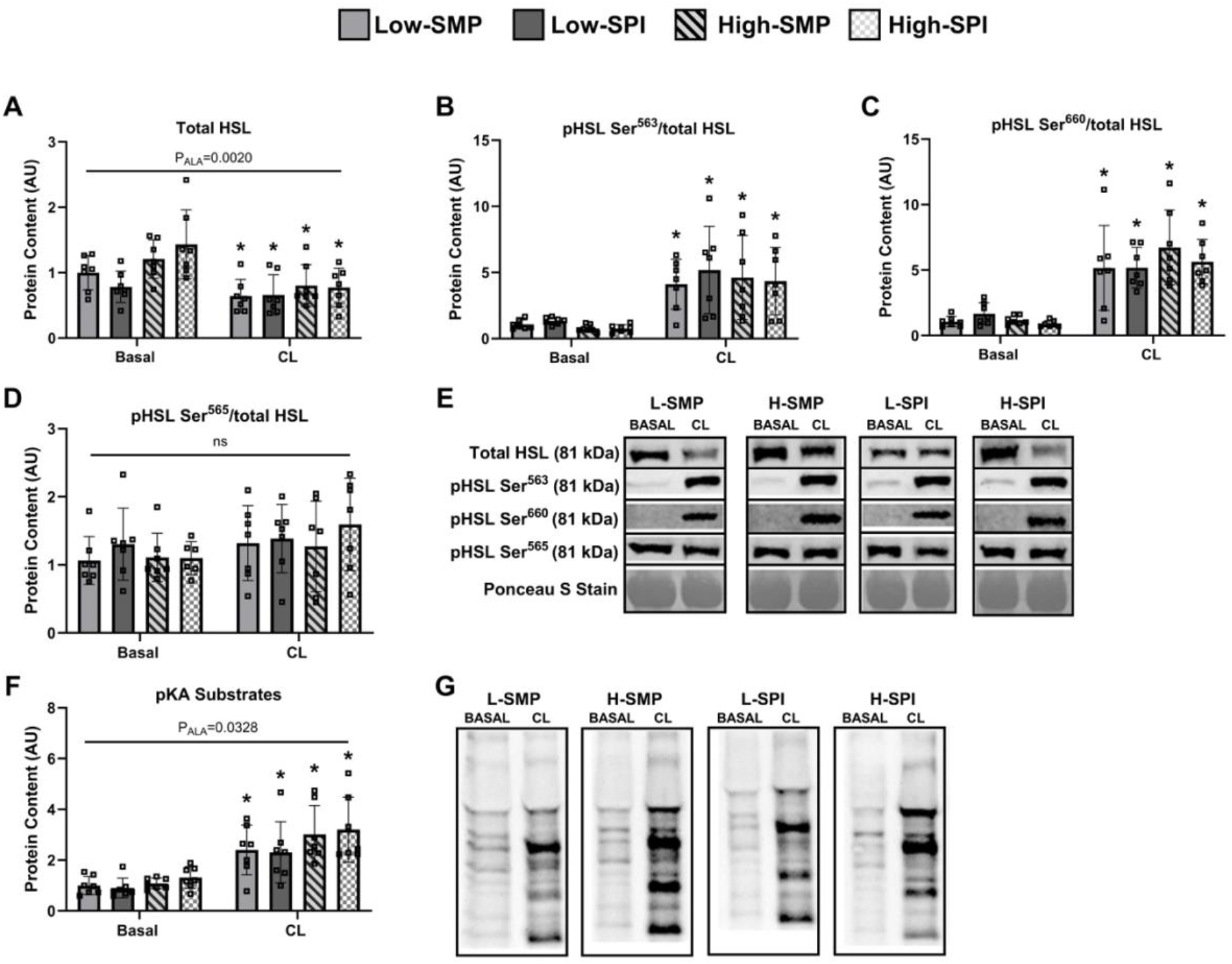
Markers of lipolysis in culture scWAT in basal and stimulated conditions. The protein abundance for total HSL (A), pHSL Ser^563^ (B), pHSL Ser^660^ (C), and pHSL Ser^565^ (D) were measured in cultured scWAT samples from mice (n=6-7 mice per group) in both basal and CL 316,243 (CL) stimulated conditions. Representative blots are provided in (E). Briefly, each primary antibody was run on an individual gel and normalized to its respective Ponceau S stain to account for any differences in sample loading. Only a single representative Ponceau S stain is shown in (E). General confirmation of β-adrenergic signaling with CL 316,243 (CL) is shown in F and corresponding representative blots in G. Data was analyzed by 2-way ANOVA and reported as mean ± standard deviation. * indicates significant differences between matching CL and basal conditions. P_ALA_, main effect of dietary ALA, ns = not significant.

**Figure 10.**
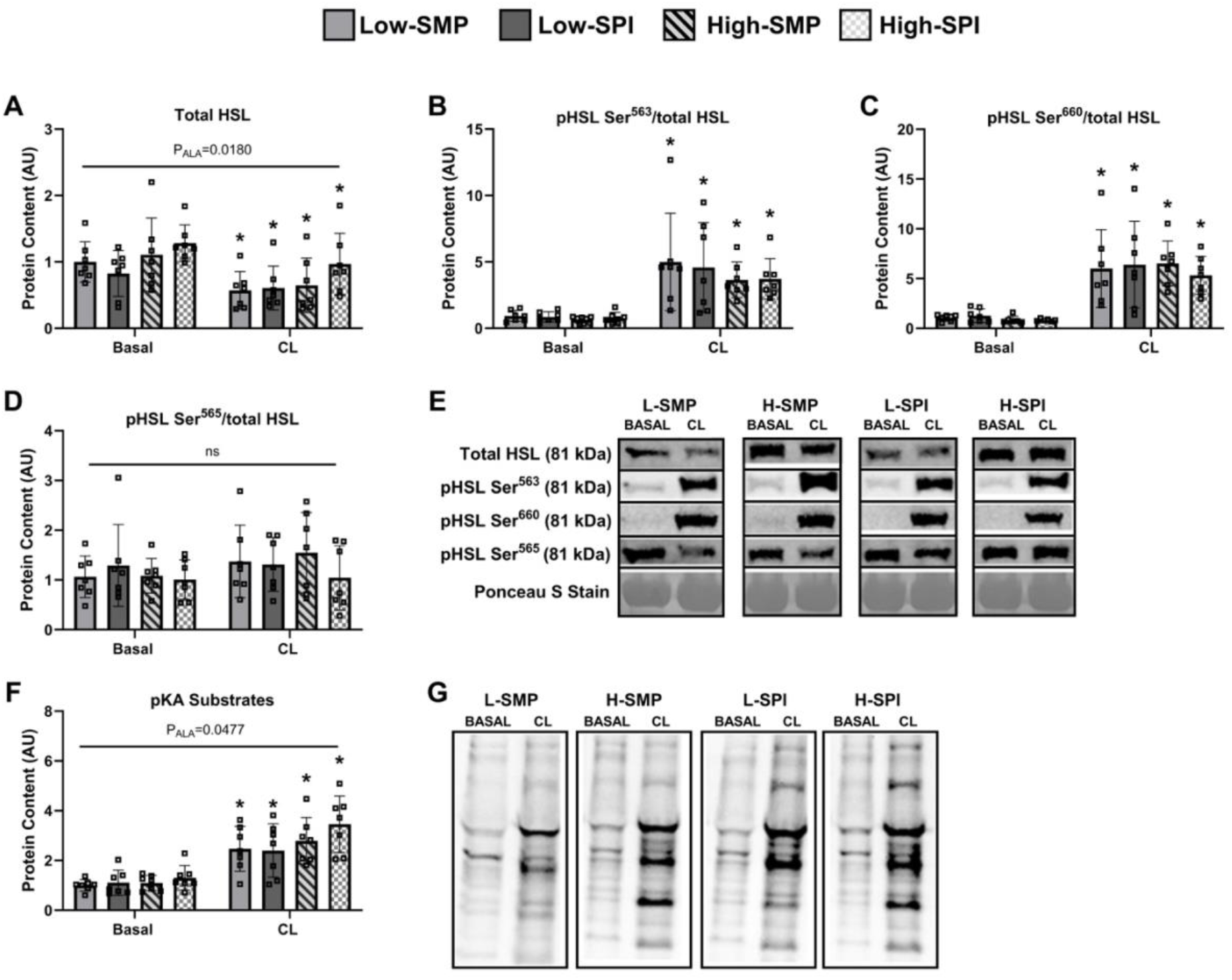
Markers of lipolysis in culture vWAT in basal and stimulated conditions. The protein abundance for total HSL (A), pHSL Ser^563^ (B), pHSL Ser^660^ (C), and pHSL Ser^565^ (D) were measured in cultured scWAT samples from mice (n=6-7 mice per group) in both basal and CL 316,243 (CL) stimulated conditions. Representative blots are provided in (E). Briefly, each primary antibody was run on an individual gel and normalized to its respective Ponceau S stain to account for any differences in sample loading. Only a single representative Ponceau S stain is shown in (E). General confirmation of β-adrenergic signaling with CL 316,243 (CL) is shown in F and corresponding representative blots in G. Data was analyzed by 2-way ANOVA and reported as mean ± standard deviation. * indicates significant differences between matching CL and basal conditions. P_ALA_, main effect of dietary ALA, ns = not significant.

## DISCUSSION

Overall, the present study revealed that female mice showed little-to-no response in markers of lipolysis following the consumption of high-ALA diets, regardless of background dietary protein. Body weight and WAT depot weights were not impacted by ALA content or dietary protein. Serum metabolic markers were unchanged with high ALA intake, while total cholesterol and the NEFA/glycerol ratio were lower in female mice fed SPI diets versus SMP diets. Alterations in lipolytic markers in response to ALA supplementation were depot-specific and, in general, rather limited. Neither ALA nor background dietary protein altered lipolytic markers in scWAT, while mice fed the high-ALA diets showed higher ATGL content in vWAT. Further, mice fed SPI diets had higher total HSL and lower pHSL-Ser565 in vWAT. Finally, ALA content caused minor differences in total HSL in cultured WAT samples, but no changes were observed in the various pHSL sites. Collectively, this work suggests that high ALA diets in female mice had minimal effects on adiposity and markers of WAT lipolysis, regardless of background dietary protein.

In contrast with findings from this study, we recently reported that WAT depots from male mice showed a more robust and consistent response to high-ALA diets, with no effect of background dietary protein [27]. Specifically, male mice consuming high-ALA diets showed higher body weight, higher WAT depot mass, as well as lower serum TAG and higher serum glycerol. Female mice did not show any differences in body and WAT depot weights in response to the same diets, or in serum TAG and glycerol levels. Male mice showed consistent changes in lipolytic markers in both WAT depots in response to high ALA diets, while female mice did not. Specifically, total ATGL was lower in both WAT depots while total HSL was higher in scWAT in male mice. In contrast, female mice did not show any changes in lipolytic markers in scWAT, and only minor differences in vWAT in response to the diets. When considered together, these two studies suggest that male mice are more sensitive to dietary ALA regulation of WAT lipolysis compared to female mice, although no direct sex comparison was performed due to insufficient statistical power.

Lipolysis is a highly regulated process that is controlled by numerous factors that are influenced by biological sex. For example, sexual dimorphism in hormonal response, adipose tissue distribution, and gene expression are documented to impact the regulation of lipolysis, thus highlighting the need to conduct nutritional studies in both males and females. Hormone signalling and sensitivity vary between males and females, particularly those controlling lipolysis such as insulin, which inhibits lipolysis, and catecholamines, which stimulate lipolysis, thus impacting metabolic outcomes and responsiveness to dietary intervention [31,32]. Females are reported to be more insulin-sensitive than males, suggesting that even at equivalent hormone levels, sex differences in the regulation of WAT lipolysis may exist [31]. Additionally, soy protein was reported to improve insulin sensitivity in male murine models compared to casein [16,33]; however, whether this extends to females is unknown, despite their increased insulin sensitivity. These past findings reinforce the importance of studying the impact of diet composition on whole-body glucose homeostasis in female mice. Similar to our previous findings in male mice [27], we found no difference in glucose (GTT) or insulin (ITT) tolerance following the different diets in female mice. These results do not appear to align with those presented in a recent meta-analysis suggesting that ALA supplementation significantly decreased serum fasting insulin levels in females compared to males [34]; however, fasting insulin levels were not measured in our studies. Consequently, additional work in females fed high ALA diets varying in background protein are warranted.

Adipose tissue depots also show sexual dimorphism, which is known to influence lipid metabolism in both basal and catecholamine-stimulated conditions. The two main WAT depots, scWAT and vWAT, have distinct metabolic roles that differentially impact overall tissue metabolism. In humans, females have proportionally more scWAT than vWAT, a distribution associated with lower basal lipolysis and increased lipid storage compared to males [32,35]. Conversely, males demonstrate increased lipolytic responsiveness and enhanced lipid mobilization with elevated vWAT content [32,35]. When comparing our previous results in male mice with those generated in the present study in female mice, we hypothesize that female mice may require increased catecholamine stimulation to activate lipolytic markers while males may show a response with less stimulation. However, we acknowledge that this notion does not appear to align with results reported in a recent human study. Indeed, Massier et al. [36] reported that while adipocytes isolated from subcutaneous adipose tissue from men had increased sensitivity to catecholamines, the maximum lipolytic effect was higher in women. Collectively, these findings suggest that the regulation of lipolysis varies between sexes, although inconsistencies across studies and species highlight the need for further investigation of sexual dimorphism.

Finally, sex-specific differences in lipolysis may be attributed to a differential responsiveness to dietary isoflavones [27]. Soy isoflavones are phytoestrogens that structurally resemble estrogen. Based on past literature, isoflavones may underlie the different lipolytic responses to diets containing casein (a dairy protein) and soy [37]. For example, Kim et al. [38] reported that a soybean embryo extract, which is a rich source of isoflavones, increased lipolysis in brown adipose tissue in male mice relative to vehicle-treated controls; however, this study did not examine WAT or female mice. Although no direct statistical comparisons were possible between our two studies in male and female mice, dietary protein appeared to produce opposing responses in lipolytic protein content in females compared to males. To the best of our knowledge, no study has directly compared lipolytic response to soy-containing diets across both sexes; however, past sex-specific studies suggest future investigations of sexual dimorphism are warranted. For example, Xiao et al. reported that female rats fed a diet containing casein showed an upregulation in hepatic lipolytic liver proteins when supplemented with soy isoflavones [39]. In contrast, Zanella et al. reported that supplementing male mice with genistein did not alter ATGL and HSL gene expression [40].

Further, the differential responsiveness to dietary isoflavones may also explain the sex-specific contrasts in serum markers [27]. It was recently demonstrated by Kishida et al. [41] that female rats provided an isoflavone-rich soybean extract exhibited dose-dependent reductions in total serum cholesterol and TAG, whereas no such response was observed in males. As such, we anticipated that the effect of background dietary protein on serum markers would be more pronounced in females than males. In our study, female mice consuming the SPI-diet showed lower serum total cholesterol and a NEFA/glycerol ratio compared to female mice fed the SMP-diet; an effect not seen in our previous study in male mice. This difference in serum cholesterol is supported by prior studies reporting that female murine models, but not males, experience reductions in serum cholesterol when consuming high-isoflavone diets compared to low-isoflavone diets [42,43]. Female mice additionally had increased serum NEFA when consuming a casein diet supplemented with isoflavones compared to male mice provided the same diet [39]. Collectively, these examples reinforce the importance of conducting studies in both male and female models to ensure the generalizability of nutritional research findings.

Limitations in the present study are similar to those outlined in our previous study using male mice [27]. For example, we did not measure monoacylglycerol lipase (MGL), which also contributes to lipolysis, or markers of FA re-esterification and lipogenesis that are relevant to WAT lipid buffering capacity. However, there are several additional limitations to consider regarding the present study in female mice. First, we did not control for estrogen levels, which may complicate our interpretation of how experimental diets regulated lipolytic markers. Females exhibit fluctuations in hormone levels, particularly estrogen, that can impact LPL-mediated TAG clearance and downregulate HSL-mediated lipolysis [35,44]. In this study, scWAT showed no response to ALA or background dietary protein, whereas vWAT had higher total HSL and lower pHSL Ser^565^ in female mice fed the SPI-diet, which may relate to estrous-related hormonal variation. Second, WAT depot distribution and adipocyte size vary by sex, but this was not measured in our studies. Since males generally exhibit greater adipocyte size, this could provide insight into why lipolytic markers were more consistently changed with high-ALA in male mice in our previous study [20,45]. This is consistent with evidence that larger adipocytes exhibit greater lipolytic activity on a per-cell basis [46]. In addition, although the experimental methodology was identical to the previous male study, these two analyses were conducted independently and direct statistical comparison between sexes was therefore not possible due to insufficient statistical power. Future studies will therefore help determine whether variation in WAT lipid metabolism relates to inherent sexual dimorphism or differential response to diet intervention.

## CONCLUSION

Our results indicate that ALA and background dietary protein have minimal effects on body weight, WAT depot mass and the regulation of lipolysis in female mice fed moderate-fat diets. Future research designs should allow for sex as a biological variable to ensure that observed responses are appropriately interpreted as either treatment responses or sex-dependent differences. Since the existing knowledge base regarding biological mechanisms and response to diets generally stem from data collected from male animal models, it is imperative that studies are conducted in female animal models to ensure the generalizability of findings in nutritional sciences. Further investigation is required to understand how ALA content, protein source, and biological sex may interact to regulate WAT lipid buffering capacity and the corresponding implications on cardiometabolic health.

## AUTHOR CONTRIBUTIONS

M.J.C conducted data analysis, data interpretation, and drafted the original manuscript. S.E.W conducted mouse and wet lab experiments and collated the data. M.G.S. and A.N.K. contributed to data collection and interpretation, and manuscript editing. F.C. conducted serum analyses and manuscript editing. D.C.W contributed to study conceptualization, funding acquisition, and manuscript editing. D.M.M. conceptualized the study design, contributed to the data analyses and interpretation, manuscript editing and acquired funding. All authors approved the final version of the manuscript.

## ACKNOWLEDGEMENTS

The authors would like to thank Chrystèle Jouve (INRA) for her technical expertise with the clinical chemistry analyzer. This research was supported by Dairy Farmers of Canada. As per the research agreement, Dairy Farmers of Canada had no role in the design and conduct of the study, data collection, and analysis or interpretation of the results as well as the decision to publish the findings. M.J.C was supported by a CBS Graduate Tuition Scholarship from the University of Guelph. S.E.W was supported by an NSERC-CGSM award. M.G.-S. was supported by a CBS-International PhD Graduate Research Assistantship and an International Doctoral Tuition Scholarship from the University of Guelph.

## CONFLICT OF INTEREST

The authors declare no conflicts of interest.

## Abbreviations

ALA: alpha-linolenic acid
ATGL: adipose triglyceride lipase
ATOC: adipose tissue organ culture
DHA: docosahexaenoic acid
EPA: eicosapentaenoic acid
FA: fatty acids
HSL: hormone-sensitive lipase
LPL: lipoprotein lipase
NEFA: non-esterified fatty acids
pKA: protein kinase A
PL: phospholipid
scWAT: subcutaneous white adipose tissue
SMP: skim milk protein
SPI: soy protein isolate
TAG: triglyceride
vWAT: visceral white adipose tissue
WAT: white adipose tissue

## Notes

### Competing Interest Statement

The authors have declared no competing interest.

## REFERENCES

[1] S.-M. An, S.-H. Cho, J. C. Yoon, Diabetes Metab. J., 2023, 47, 595–611.

[2] K. N. Frayn, Diabetologia, 2002, 45, 1201–1210.

[3] A. Sakers, M. K. De Siqueira, P. Seale, C. J. Villanueva, Cell 2022, 185, 419–446.

[4] A. E. Carrillo, M. Vliora, Nutrients 2023, 15, 4811.

[5] F. Haugen, C. A. Drevon, Proc. Nutr. Soc. 2007, 66, 171–182.

[6] A. Fernández-Quintela, I. Churruca, M. P. Portillo, Public Health Nutr. 2007, 10, 1126– 1131.

[7] L. Madsen, L. S. Myrmel, E. Fjære, J. Øyen, K. Kristiansen, Front. Physiol. 2018, 9, 1792.

[8] F. Shahidi, P. Ambigaipalan, Annu. Rev. Food Sci. Technol. 2018, 9, 345–381.

[9] C. C. Tai, S. T. Ding, J. Nutr. Biochem. 2010, 21, 357–363.

[10] S. Lorente-Cebrián, A. G. V. Costa, S. Navas-Carretero, M. Zabala, J. A. Martínez, M. J. Moreno-Aliaga, J. Physiol. Biochem. 2013, 69, 633–651.

[11] G. C. Shearer, O. V. Savinova, W. S. Harris, Biochim. Biophys. Acta BBA - Mol. Cell Biol. Lipids 2012, 1821, 843–851.

[12] C. Wang, B. Hucik, O. Sarr, L. H. Brown, K. R. D. Wells, K. R. Brunt, M. T. Nakamura, E. Harasim-Symbor, A. Chabowski, D. M. Mutch, J. Lipid Res. 2023, 64, 100376.

[13] B. MacLeod, C. Wang, L. H. Brown, E. Borkowski, M. T. Nakamura, K. R. Wells, K. R. Brunt, E. Harasim-Symbor, A. Chabowski, D. M. Mutch, J. Lipid Res. 2024, 65, 100642.

[14] J. N. Smorenburg, K. Hodun, P. V. McTavish, C. Wang, M. A. Pinheiro, K. R. D. Wells, K. R. Brunt, M. T. Nakamura, A. Chabowski, D. M. Mutch, Mol. Nutr. Food Res. 2025, 69, e202400721.

[15] T. L. Blasbalg, J. R. Hibbeln, C. E. Ramsden, S. F. Majchrzak, R. R. Rawlings, Am. J. Clin. Nutr. 2011, 93, 950–962.

[16] C. Ascencio, N. Torres, F. Isoard-Acosta, F. J. Gómez-Pérez, R. Hernández-Pando, A. R. Tovar, J. Nutr. 2004, 134, 522–529.

[17] L. Noriega-López, A. R. Tovar, M. Gonzalez-Granillo, R. Hernández-Pando, B. Escalante, P. Santillán-Doherty, N. Torres, J. Biol. Chem. 2007, 282, 20657–20666.

[18] J. W. Anderson, B. M. Johnstone, M. E. Cook-Newell, N. Engl. J. Med. 1995, 333, 276– 282.

[19] P. Kaur, R. Kaur, S. Sharma, S. Kaur, Crit. Rev. Food Sci. Nutr. 2025, 1–17.

[20] E. Fuente-Martín, P. Argente-Arizón, P. Ros, J. Argente, J. A. Chowen, Adipocyte 2013, 2, 128–134.

[21] N. Boulet, A. Briot, J. Galitzky, A. Bouloumié, Biomedicines 2022, 10, 2615.

[22] D. N. Costa, S. Santosa, M. D. Jensen, Physiol. Rev. 2025, 105, 897–934.

[23] B. T. Palmisano, L. Zhu, R. H. Eckel, J. M. Stafford, Mol. Metab. 2018, 15, 45–55.

[24] O. Varlamov, C. L. Bethea, C. T. Roberts, Front. Endocrinol. 2014, 5, 241.

[25] C. Koutsari, R. Basu, R. A. Rizza, K. S. Nair, S. Khosla, M. D. Jensen, J. Clin. Endocrinol. Metab. 2011, 96, 541–547.

[26] E. Blaak, Curr. Opin. Clin. Nutr. Metab. Care 2001, 4, 499–502.

[27] S. E. Woods, M. Gonzalez-Soto, A. N. King, F. Capel, D. C. Wright, D. M. Mutch, J. Nutr. Biochem. 2025, 110185.

[28] P. Arner, Ann. Med. 1995, 27, 435–438.

[29] P. M. Kris-Etherton, J. A. Grieger, T. D. Etherton, Prostaglandins Leukot. Essent. Fatty Acids 2009, 81, 99–104.

[30] K. J. Hintze, A. D. Benninghoff, C. E. Cho, R. E. Ward, Adv. Nutr. Bethesda Md 2018, 9, 263–271.

[31] M. Keuper, M. Jastroch, Mol. Cell. Endocrinol. 2021, 533, 111337.

[32] S. Wang, K. G. Soni, M. Semache, S. Casavant, M. Fortier, L. Pan, G. A. Mitchell, Mol. Genet. Metab. 2008, 95, 117–126.

[33] A. Ji, W. Chen, C. Liu, T. Zhang, R. Shi, X. Wang, H. Xu, D. Li, Food Funct. 2023, 14, 5752–5767.

[34] S. Mohammadi, D. Ashtary-Larky, N. Alaghemand, I. Alavi, M. Erfanian-Salim, P. S. Pirayvatlou, Y. Ettehad, A. Borzabadi, M. Mehrbod, O. Asbaghi, Nutr. Metab. Cardiovasc. Dis. NMCD 2026, 36, 104370.

[35] V. Xega, J.-L. Liu, Med. Rev. 2021 2024, 4, 284–300.

[36] L. Massier, D. P. Andersson, N. Viguerie, J. Zhong, D. Zareifi, A. G. Kerr, D. Langin, P. Arner, iScience 2025, 28, 113988.

[37] M. Gonzalez-Soto, D. M. Mutch, Adv. Nutr. Bethesda Md 2021, 12, 980–994.

[38] M. Kim, S. Im, Y. K. Cho, C. Choi, Y. Son, D. Kwon, Y.-S. Jung, Y.-H. Lee, Biomolecules 2020, 10, 1394.

[39] C. W. Xiao, C. M. Wood, D. Weber, S. A. Aziz, R. Mehta, P. Griffin, K. A. Cockell, Genes Nutr. 2014, 9, 373.

[40] I. Zanella, E. Marrazzo, G. Biasiotto, M. Penza, A. Romani, P. Vignolini, L. Caimi, D. Di Lorenzo, Eur. J. Nutr. 2015, 54, 1095–1107.

[41] T. Kishida, T. Mizushige, M. Nagamoto, Y. Ohtsu, T. Izumi, A. Obata, K. Ebihara, Biosci. Biotechnol. Biochem. 2006, 70, 1547–1556.

[42] O. Mezei, Y. Li, E. Mullen, J. S. Ross-Viola, N. F. Shay, Physiol. Genomics 2006, 26, 8– 14.

[43] O. Mezei, W. J. Banz, R. W. Steger, M. R. Peluso, T. A. Winters, N. Shay, J. Nutr. 2003, 133, 1238–1243.

[44] G. C. Burdge, S. A. Wootton, Br. J. Nutr. 2002, 88, 411–420.

[45] P. Löfgren, J. Hoffstedt, M. Rydén, A. Thörne, C. Holm, H. Wahrenberg, P. Arner, J. Clin. Endocrinol. Metab. 2002, 87, 764–771.

[46] J. Laurencikiene, T. Skurk, A. Kulyté, P. Hedén, G. Aström, E. Sjölin, M. Rydén, H. Hauner, P. Arner, J. Clin. Endocrinol. Metab. 2011, 96, E2045–2049.

